# Effects of variability in manually contoured spinal cord masks on fMRI co-registration and interpretation

**DOI:** 10.1101/2022.03.25.485810

**Authors:** Mark A. Hoggarth, Max C. Wang, Kimberly J. Hemmerling, Andrew D. Vigotsky, Zachary A. Smith, Todd B. Parrish, Kenneth A. Weber, Molly G. Bright

**Affiliations:** Department of Physical Therapy and Human Movement Sciences, Feinberg School of Medicine, Northwestern University, Chicago, IL, USA; Department of Biomedical Engineering, McCormick School of Engineering, Northwestern University, Evanston, IL, USA; Department of Statistics, Weinberg College of Arts and Sciences, Northwestern University, Evanston, IL, USA; Department of Neurological Surgery, University of Oklahoma Health Sciences Center, Oklahoma City, OK, USA; Department of Radiology, Feinberg School of Medicine, Northwestern University, Chicago, IL, USA; Systems Neuroscience and Pain Lab, Department of Anesthesiology, Perioperative and Pain Medicine, Stanford University, Palo Alto, CA, USA

**Author notes:** <u>Corresponding Author:</u> Molly G. Bright, D.Phil., 645 N. Michigan Ave. Suite 1100, Chicago, IL 60611. Joint First Authors (Mark A. Hoggarth, Max C. Wang).

**Keywords:** Functional MRI, Spinal cord, Contouring, Image Registration, Reliability

## Abstract

Functional magnetic resonance imaging (fMRI) of the human spinal cord (SC) is a unique non-invasive method for characterizing neurovascular responses to stimuli. Group-analysis of SC fMRI data involves co-registration of subject-level data to standard space, which requires manual masking of the cord and may result in bias of group-level SC fMRI results. To test this, we examined variability in SC masks drawn in fMRI data from 21 healthy participants from a completed study mapping responses to sensory stimuli of the C7 dermatome. Masks were drawn on temporal mean functional image by eight raters with varying levels of neuroimaging experience, and the rater from the original study acted as a reference. Spatial agreement between rater and reference masks was measured using the Dice Similarity Coefficient, and the influence of rater and dataset was examined using ANOVA. Each rater’s masks were used to register functional data to the PAM50 template. Gray matter-white matter signal contrast of registered functional data was used to evaluate the spatial normalization accuracy across raters. Subject- and group-level analyses of activation during left- and right-sided sensory stimuli were performed for each rater’s co-registered data. Agreement with the reference SC mask was associated with both rater (F(7,140) = 32.12, P < 2×10^−16^, *η*^2^ = 0.29) and dataset (F(20,140) = 20.58, P < 2×10^−16^, *η*^2^ = 0.53). Dataset variations may reflect image quality metrics: the ratio between the signal intensity of spinal cord voxels and surrounding cerebrospinal fluid was correlated with DSC results (p<0.001). As predicted, variability in the manually-drawn masks influenced spatial normalization, and GM:WM contrast in the registered data showed significant effects of rater and dataset (rater: *F*(8,160) = 23.57, *P* < 2×10^−16^, *η*^2^ = 0.24; dataset: *F*(20,160) = 22.00, *P* < 2×10^−16^, *η*^2^ = 0.56). Registration differences propagated into subject-level activation maps which showed rater-dependent agreement with the reference. Although group-level activation maps differed between raters, no systematic bias was identified. Increasing consistency in manual contouring of spinal cord fMRI data improved co-registration and inter-rater agreement in activation mapping, however our results suggest that improvements in image acquisition and post-processing are also critical to address.

## Introduction

Functional magnetic resonance imaging (fMRI) of the spinal cord is a technique for understanding neurovascular responses to sensory and motor stimuli.[1-3] fMRI studies of the cord have demonstrated neural activation correlates to physiological mechanisms of sensation, illustrating dermatomal patterns[4-8], and motor control laterality.[9] Furthermore, studies have demonstrated interactions between the cortex and spinal cord in both sensory and motor learning paradigms that further our understanding of human neurology.[10-11] Encouraged by observations of coordinated intrinsic activity within functional brain networks, researchers have successfully demonstrated such “connectivity” properties within the spinal cord, and between the cord and brain.[11-13] Uniquely positioned to provide noninvasive functional mapping of large segments of the spinal cord in humans, spinal cord fMRI is poised to play a critical role in understanding both typical and pathologic sensation and movement.

However, while significant advances have been made over the years, the breadth of literature studying the spinal cord still lags behind that of brain fMRI research, in part due to remaining technical challenges, including those related to cord anatomy. Specifically, analysis and interpretation of spinal cord fMRI data is hindered by the large changes in magnetic susceptibility of tissues approximating the cord, the small size of the target neural tissues (typically leading to low signal-to-noise ratio), and physiological noise from cardiac and respiratory processes.[14-18] Several analytical tools have been developed to improve characterization of spinal cord fMRI data. These tools include, but are not limited to the Spinal Cord Toolbox, the Neptune Toolbox, and Pantheon (formerly spinalfMRI8).[19-20] There are also specific analysis strategies developed for spinal cord fMRI denoising such as physiological noise modeling[17,21], slice-wise motion correction[19-20], anisotropic spatial smoothing [19], and principal and independent component analysis based denoising[22-24].

At present, these noise-reduction strategies can dramatically improve the quality of fMRI data. However, in typical 3T fMRI scans, it remains challenging to confidently interpret activation maps in individual subject data. As in brain fMRI, an established way to make statistical inferences across the sample or a population is to combine fMRI data from many individuals for group-level analysis. One way to do this is by using region of interest (ROI)-based group analysis relying on regions defined at the subject level. However, in order to retain complete, voxelwise spatial information, it is often desirable to generate group-level activation maps. This process necessitates the co-registration of fMRI data to a common or standard space to facilitate comparisons across subjects with variable cord anatomy.[25] To this end, standard spinal cord templates and techniques for registration have been developed.[18] Notably, and unlike brain fMRI co-registration, nearly all spinal cord fMRI registration techniques require user input. One early method that was iteratively developed required the user to indicate various reference lines to inform affine transformation to a reference image with minimal curvature.[26-29] This technique was applied on sagittally acquired slices, and in 2015 an automated version was published.[30] Around the same time, different techniques were developed for axially acquired data. One study performed normalization by manually identifying the center of the cord in each slice, then performing an in-plane translation (2 degrees of freedom) to match a reference.[11] A similar approach using a manually defined spinal cord mask and 4 degrees of freedom (translation and scaling) was also developed.[6] Building on these axial normalization techniques, the sct_register_multimodal function was introduced as part of the spinal cord toolbox (SCT) in 2017. This was the first nonlinear registration algorithm for spinal cord fMRI and it has since been used in various studies of the cord.[7, 9, 19, 31]

When using sct_register_multimodal to register functional data to a template, it is advisable to use the warping field from a previously completed structural to template registration to initialize the algorithm and exploit the high-resolution information available for the individual subject. Additionally, due to the challenges of image distortion in EPI data, it is recommended for this function that the user inputs a binary spinal cord mask in native fMRI space to inform registration. It is common for this input mask to be manually defined, as there are currently no reliable algorithms for segmenting the spinal cord in functional data. Note that spinal cord data acquired with non-EPI methods, and with high-resolution data acquired from a common field-of-view, may facilitate alternative automated approaches for cord masking that would not be impacted by the manual masking element explored in this paper. The remaining challenges of spinal cord fMRI data quality (e.g., low signal-to-noise, residual physiologic and motion artifacts, and poor contrast between spinal cord tissue and the surrounding cerebrospinal fluid (CSF)) may lead to subjective differences when contouring the cord. These differences could result in systematic bias in data co-registration and thus alter individual and group analysis results. The extent of this source of variability in spinal cord fMRI processing pipelines is relatively unknown.

Standardization of both imaging protocols and processing pipelines has been recommended to improve the robustness of spinal cord fMRI findings.[15, 18] In brain fMRI, differences in functional image processing pipelines have been shown to have large potential impacts on the resultant findings of a study.[32] Bowring et al. observed that variability in results from task fMRI in the brain are heterogeneous, depending on the input dataset and potentially each aspect of the image processing pipeline (including registration).[32] Furthermore, study results were also impacted by the software package utilized.[32] While there are multiple software packages for spatial normalization of fMRI in the brain[33-35], spinal cord studies have fewer options or must create bespoke techniques for registration to labeled structural templates of the cord.[18-19, 36] Given the lack of a unified method of preprocessing spinal cord fMRI data, reproducibility of reported results is likely to be limited, and improvements are needed to standardize image processing and reduce sources of variability and bias at every stage of analysis.

In this work, we specifically assess the impact of variable manual contouring of the spinal cord in native fMRI image space, as needed for spatial normalization of individual fMRI datasets to a standard template space. To achieve this, we examine the effects of mask variability at different stages of a single analysis pipeline for spinal cord fMRI using a previously published study dataset. We characterize the variability in spinal cord masks achieved by eight raters with varying levels of image analysis experience, with respect to a “reference” rater from the original study and publication. We then demonstrate how this variability is propagated following registration of functional imaging data to a standard spinal cord template image. We subsequently run individual- and group-level analyses for a sensory stimulus task, using each rater’s masks during co-registration, to assess the impact on fMRI activation patterns at the single-subject and overall study level. Finally, we discuss the causes of this variability, their importance, and make recommendations for prioritizing future improvements in spinal cord fMRI.

## Methods

### 2.1 Image acquisition and experimental protocol

This work utilized a subset of anatomical and functional MRI data from 24 healthy participants from a previous study.[37] Images were acquired using a 3T Siemens Prisma scanner (Siemens, Erlangen, Germany), utilizing a 64-channel head/neck coil and a SatPad™ cervical collar (SatPad Clinical Imaging Solutions, West Chester, PA, USA). Anatomical T2-weighted images were acquired covering the cervical and upper thoracic spine, using the SPACE sequence (Siemens, Erlangen, Germany), with parameters: TR = 1500ms, TEeff = 135 ms, echo train = 74, flip angle = 90°/140°, slices = 64, effective voxel size = 0.8 × 0.8 × 0.8 mm^3^, iPAT acceleration factor = 3, interpolated in-plane resolution = 0.4 × 0.4 mm^2^.[38-39] T2*-weighted functional scans of the cervical spinal cord were acquired, with 25 axially acquired slices centered at the C5 vertebral level, using a gradient-echo echo-planar-imaging sequence with ZOOMit selective field-of-view.[40-42] Functional imaging parameters were: TR3D = 2000 ms, TE = 30 ms, flip angle = 80°, volumes = 450, slice order was interleaved, field-of-view = 128 × 44 mm^2^, acquisition matrix = 128 × 44 voxels, in-plane resolution = 1 × 1 mm^2^, slice thickness = 3 mm, two dummy volumes discarded.

During each functional scan, alternating left and right tactile stimuli were applied to the dorsum of each hand in the C7 dermatomal region.[37] Stimulation was applied manually at approximately 2 Hz by examiners in the scan room. Stimulation lasted for 15 sec on each hand and was interspersed with a 15 second rest period.

Preprocessing of the functional MRI time series was performed as in the original study.[11] Motion correction was performed in two phases using FMRIB’s Linear Registration Tool (FLIRT) for 3D rigid body alignment (6 degrees of freedom), followed by rigid 2D slice-wise (axial) alignment (2 degrees of freedom; x- and y-translation only) [34, 43], and was optimized for a binary masked region around the spinal canal. Temporal mean images were calculated from these preprocessed functional data. fMRI data were further denoised by removing periodic physiological noise confounds using the PNM (Physiological Noise Modelling) tool in FSL [21, 44], warped to the PAM 50 spinal cord template [45], and then smoothed with a 2mm^3^ full width half maximum Gaussian smoothing kernel prior to individual- and group-level activation mapping.[46]

### 2.2 Manual contouring

Raters (N=8) with differing levels of functional neuroimaging experience were recruited to manually contour the spinal cord on 21 temporal mean functional images, after 3 of the 24 datasets were utilized for rater training. Additionally, as this dataset was collected for a previously completed and published study,[37] the original masks from that work were included as “reference” contours. The reference masks (REF) are provided by a researcher with 10 years of experience with spinal cord fMRI (KAW). Including the reference contours, a total of 9 sets of masks were used in subsequent analyses. A description of each rater’s background and experience in the neuroimaging field is given in Table 1. Note that while raters with no previous experience would not be expected to generate masks in a typical preprocessing pipeline, their inclusion in this study serves two purposes. First, we use these raters to establish a baseline level of performance to which more experienced raters can be compared. Second, by considering the most extreme variability within reason, we leave no room to doubt whether our range of raters is wide enough to observe an effect.

**Table 1.** Description of each rater’s background

| **Rater** | **Job Title** | **MRI**<br>**experience**<br>**(years)** | **SC MRI**<br>**experience**<br>**(years)** | **SC fMRI**<br>**experience**<br>**(years)** | **Research keywords** |
| --- | --- | --- | --- | --- | --- |
| Reference | Instructor | 12 | 10 | 10 | Brain, spinal cord, pain,<br>musculoskeletal MRI,<br>neurological injury |
| A | Assistant<br>professor | 14 | 2 | 2 | Cerebrovascular MRI,<br>fMRI denoising,<br>physiologic modeling |
| B | PhD student | 1 | 1 | 1 | Spinal cord fMRI,<br>analysis methods |
| C | Postdoctoral<br>researcher | 8 | 8 | 1 | Spinal Cord MRI, pain,<br>traumatic injury |
| D | Undergraduate<br>student | 1 | 1 | 1 | Spinal cord fMRI,<br>analysis methods |
| E | Undergraduate<br>student | 0 | 0 | 0 | N/A |
| F | Postdoctoral<br>researcher | 7 | 0.5 | 0.5 | Neuroscience,<br>neuroimaging,<br>cerebrovascular MRI |
| G | Undergraduate<br>student | 0 | 0 | 0 | N/A |
| H | Undergraduate<br>student | 0 | 0 | 0 | N/A |

Contouring was performed by all raters in FSLeyes.[34] During an initial training session, raters who were unfamiliar with the process of contouring (E, G, H) were guided with specific instructions on using the software to optimize image brightness, contrast, image orientation, and zoom in order to discern the spinal cord boundary. All raters were instructed to inform their contouring decisions primarily from the axial view but were also allowed to assess the continuity of their masks in the sagittal and frontal views. In addition, raters were instructed to take approximately 15 minutes per dataset. Note that this training was primarily focused on utilizing software with minimal guidance on interpreting the spinal cord boundary, so we do not expect systematic similarity in mask variability as a result of training. Following the initial training session, 3 training datasets were released to all raters to ensure competency with the masking process. Then, the remaining 21 datasets were released in 3 blocks (7 datasets each) with a randomized order for each rater. Raters were given 2 weeks to complete each block of masks.

### 2.3 Registration to standard space

Image registration was performed with the Spinal Cord Toolbox (version 4.3).[19] First, the sct_deepseg_SC function was implemented to automatically identify the cord in the high-resolution T2-weighted anatomical images[47], and the C3 and C7 vertebrae were manually labeled. The spinal cord segmentation and vertebral level labels were then used to register the anatomical image to the PAM50 spinal cord template image using the sct_register_to_template function.[19,45] The anatomical-to-template registration was performed once for each dataset and did not vary between raters. The temporal mean functional images were registered to the PAM50 template using the sct_register_multimodal function, utilizing each rater’s manually contoured masks of the cord and initial warping field (generated from the anatomical registration above) as inputs. The input rater-drawn spinal cord masks were only considered in the second step of the command (see Appendix A). In this step, voxels within the mask were heavily weighted in the warping field calculation, but added spatial regularization included in the selected algorithm (‘bsplinesyn’) warped voxels outside of the mask as well. An overview of the masking and registration process is shown in **Figure 1**.

**Figure 1:**
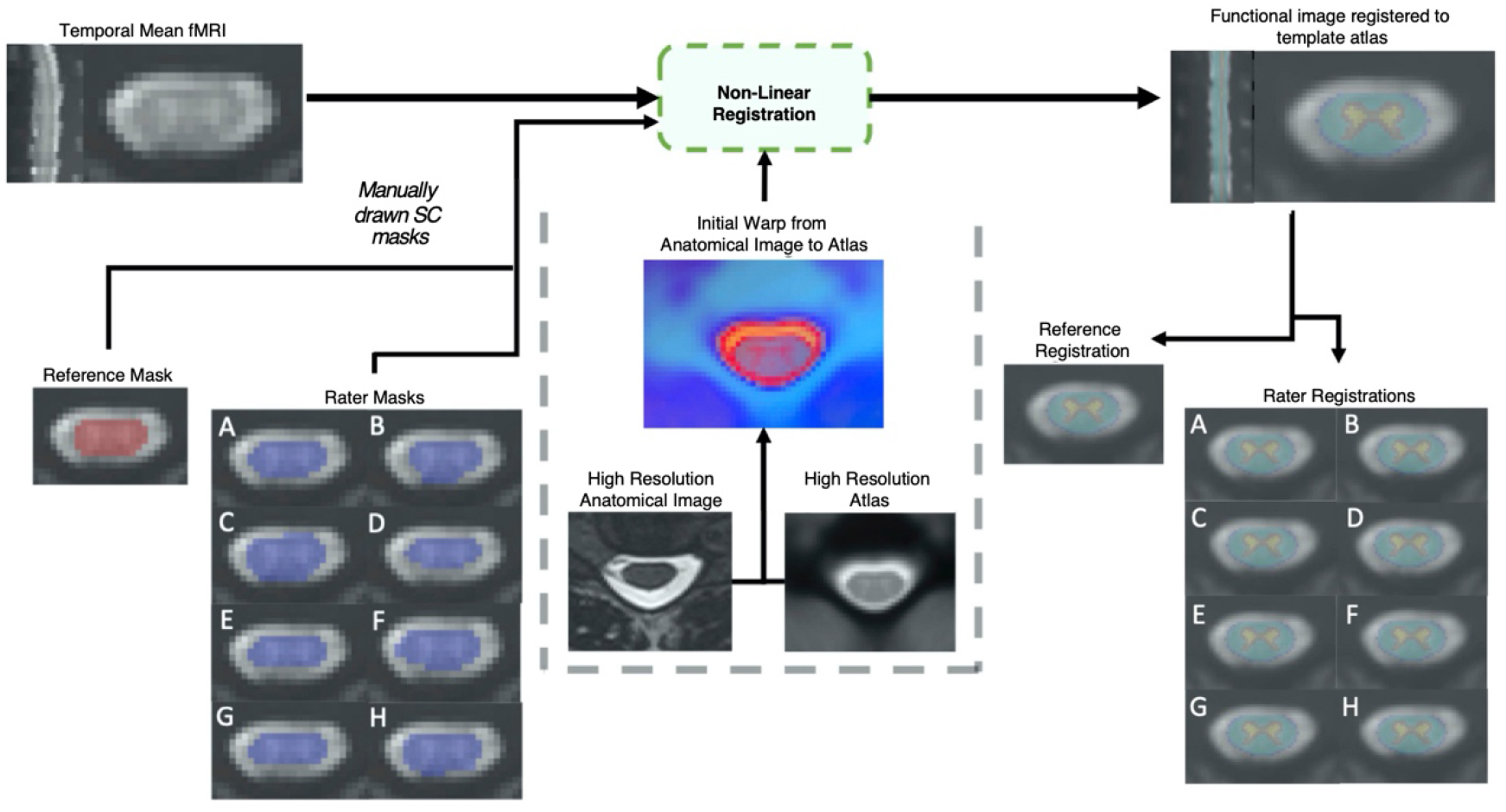
Outline of non-linear registration. A mean temporal image was created; then a reference rater and 8 raters with varied levels of experience contoured the spinal cord manually. Non-linear registration was performed with the Spinal Cord Toolbox, utilizing additional information from a high resolution anatomical T2 weighted image. The impact of the resulting registration of the functional image to template space was analyzed.

### 2.4 Variability in pre- and post-registration masks

The variability in rater-contoured masks was assessed by comparing each rater mask (RM) to the reference mask (REF). To quantify differences, the Dice Similarity Coefficient (DSC) was calculated for the total volume and for each axial slice as:

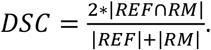

Since DSC is a proportion on the unit interval (e.g., [0,1]), it was logit-transformed for all analyses. This allows DSC to be more compatible with linear models, which are not constrained to the unit interval. Empirically, this resulted in improved model fits relative to modeling raw DSCs. ANOVA was used to compare DSC, averaged by dataset, between raters and the reference. The most superior and inferior slices were not included in these analyses due to poor image quality.

To assess the effect of fMRI image quality on rater agreement, two metrics were calculated for each dataset on the temporal mean functional image: 1) the coefficient of variation (CV) of voxels adjacent to the reference spinal cord mask and 2) the ratio between the mean signal of adjacent voxels and the mean signal within the reference spinal cord mask (Adjacent:SC ratio). For both metrics, adjacent voxels were defined by a 2-voxel dilation from the reference SC mask. The CV of adjacent voxels captures inconsistencies in CSF signal and the Adjacent:SC ratio represents the signal contrast required to contour spinal cord boundaries. A higher CV or lower Adjacent:SC ratio could obscure the delineation between spinal cord and surrounding CSF and would increase the difficulty of contouring the cord, potentially leading to increased variability in rater masks.

We compared DSC with the two metrics of image quality (CV and Adjacent:SC ratio) for each axial slice of each dataset, rather than using a summary metric for the entire volume of each dataset. When raters contoured the spinal cord images, the cord was primarily viewed in the axial plane. In this orientation, image quality can vary across slice acquisitions: for example, there is decreased efficiency of head/neck coils in more inferior slices, where magnetic field inhomogeneities may also be increased due to magnetic susceptibility variation during respiration.[48] While shimming of the magnetic field can reduce static field inhomogeneities, present methods are still incapable of fully compensating for smaller field variations due to changes in anatomical structures such as the borders between spinal discs and vertebral bodies[14-15,18], and custom dynamic shimming techniques are not routinely available to fully mitigate dynamic effects of respiration Additionally, under normal breathing conditions, the cervical spinal cord moves most in the C4-T1 region.[49-50] Combined, these factors can create significant variation in the image properties across axial slices of a given data set. Thus, we correlated slicewise metrics of image quality with DSC values for each RM (with respect to the REF) for all datasets. To do so, we calculated a separate Spearman’s *ρ* for each RM and dataset, with the variance within each RM-dataset pair arising from differences between slices—this approach prevents the dataset effects from dominating the correlations. The resulting correlations were then converted to Fisher’s *z* and averaged across datasets (within each RM). *P*-values were calculated using 5,000 max-T (or min-P) permutations, which control for multiple comparisons while accounting for covariation between outcomes. Within each permutation, we (1) scrambled the DSC values across slices within each RM-dataset pair, which assumes that the slices were exchangeable and independent, (2) calculated the Spearman’s *ρ* for each RM-dataset pair, (3) converted Spearman’s *ρ* to Fisher’s *z*, (4) averaged across datasets (within each RM), and (5) added the max absolute Fisher’s *z* to the permutation distribution. The observed absolute Fisher’s *z* values were compared to this distribution to calculate two-tailed *P*-values that were adjusted for multiple comparisons.

To evaluate the accuracy of the alignment of functional data to the standard PAM50 template space, we considered the signal contrast between gray matter (GM) and white matter (WM) voxels using regions defined by the PAM50 spinal cord atlas. Inherent signal contrast between GM and WM tissues within the spinal cord will vary across acquisitions, and the functional BOLD-weighted sequence was not designed to optimize GM:WM contrast; therefore, this value is likely small and highly variable across datasets. However, we use this metric to evaluate the relative accuracy of the rater-specific spatial normalization procedures for a given dataset: perfect alignment with the PAM50 template should lead to maximal GM:WM contrast using regions defined by that template atlas. Imperfect alignment with the PAM50 template would cause mixing of GM and WM signals across the two atlas regions, reducing the observed GM:WM contrast for that dataset. The GM:WM ratio was calculated for each dataset, following spatial normalization procedures using each rater’s spinal cord mask (or the reference). Since GM:WM is a ratio, it was first log-transformed and then fitted via an ANOVA with rater and dataset as independent variables, thus allowing us to distinguish the impact of rater on spatial normalization accuracy from inherent variability in signal contrast across the datasets. The log-transformation improved the normality of the residuals.

### 2.5 Variability in statistical activation in participant- and group-level analyses

As described in the original study, participant-level analyses were performed to characterize significant activation associated with the sensory stimuli.[37] Note that while participant-level analysis is often performed in native space, where the impact of spinal cord masks would be negligible, the analysis in the original study was done in PAM50 space to facilitate comparison across subjects and interpretation of participant-level activation in standardized coordinates. In this scheme, manual contours inform registration to template space before the GLM is applied and can therefore lead to spatial variation in activation patterns. In PAM50 space, trialwise left- and right-sided stimuli were convolved with hemodynamic response function and analyses were performed via a generalized linear model, using FILM with prewhitening.[51] Voxels with a Z-score > 2.3 (p < 0.01, uncorrected) were classified as active. These analyses were repeated for each rater. Participant-level activation maps (z-statistics) for left- and right-sided sensory stimuli were calculated using a fixed-effects analysis for each dataset and rater. Based on spinal cord anatomy, activation from the tactile stimulus was expected to localize to the ipsilateral hemicord around the C7 spinal level. We therefore initially focused our attention on ipsilateral activations, only. The spinal cord was divided into the left and right hemicord, excluding from analysis the center column of spinal cord voxels where the hemicords meet. We considered activation with the left-sided stimulus in the left hemicord, and activation with the right-sided stimulus in the right hemicord, calculating the spatial correlation (Fisher’s z) between the ipsilateral activation patterns for each rater and the reference for each individual dataset.

Group-level activation results were achieved using the participant-level activation maps derived from each rater and the reference. As in the original study, all group-level analyses were performed in the region of intersection of the functional images, again using a fixed-level analysis in FILM, where voxels with a Z-score > 2.3 and multiple comparisons correction cluster significance threshold of p < 0.05.[51] The original study found that activity was somewhat lateralized to the ipsilateral cord, but also deviated from these expectations, identifying activity more broadly distributed throughout the dorsal and ventral aspects of the cord and also superiorly and inferiorly to the expected cervical level.[37] Thus, considering the left and right hemicord regions separately, we assessed the distribution of z-statistics associated with ipsilateral activation (left cord, left stimulation; right cord, right stimulation) and contralateral activation (left cord, right stimulation; right cord, left stimulation) in the group-level results. These distributions were compared between the reference and the raters by standardizing (i.e., *z*-scoring) each of the reference’s *z*-statistics (*yi*) relative to the raters’ distribution (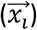) for each voxel *i* = 1, …, *n*, and then averaged the results:

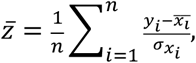

Similarly, the standard deviation was also calculated:

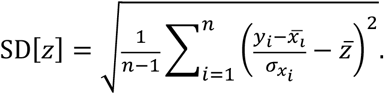

## Results

DSC comparisons between individual raters and the reference demonstrated variability across both rater and dataset as shown in **Figure 2** (rater: F(7,140) = 32.12, P < 2×10^−16^, *η*^2^ = 0.29; dataset: F(20,140) = 20.58, P < 2×10^−16^, *η*^2^ = 0.53). Note that both the rater and dataset axes in **Figure 2** are sorted by average logit-transformed DSC, and this ordering is also reflected in **Table 1** detailing rater experience (i.e. rater A achieved the highest average DSC while rater H achieved the worst). Consistent with expectation, the rater with the most MRI research experience (A) achieved the highest average DSC while 2 novice raters with no prior experience (G, H) achieved the lowest. However, it is notable that a researcher with 7 years of MRI experience (F) achieved lower DSC than a third novice rater (E). The DSC of all rater masks compared to the reference are additionally visualized by box-and-whisker plots in **Supplementary Figure 1**.

**Figure 2:**
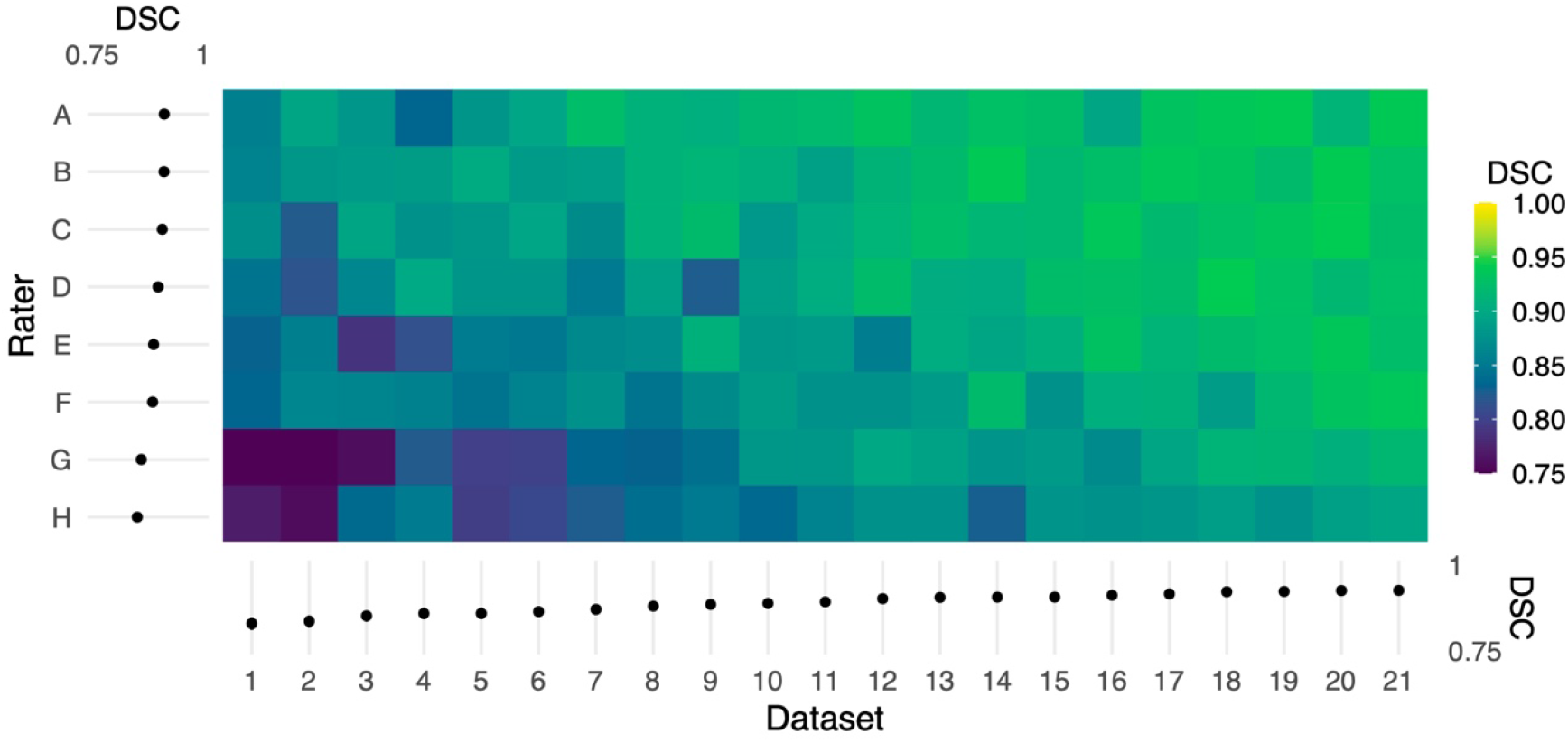
Color map of Dice similarity coefficients (DSC) comparing each rater mask with the reference mask for each of 21 datasets. Both the rater and dataset axes are organized from lowest to highest average logit-transformed DSC. The margins (left and bottom) contain estimated marginal means for each rater and dataset, and error bars indicate 95% CIs. Horizontal and vertical gradient trends indicate the effect of rater and dataset, respectively, on agreement with the reference.

The horizontal trend in **Figure 2** illustrates differences in DSC attributable to dataset features, potentially including image quality or subject anatomy. Reported in **Table 2**, the CV of voxels adjacent to the spinal cord was poorly to moderately negatively correlated with DSC for 6 raters (*ρ* = −0.50 to −0.14). The Adjacent:SC signal ratio was poorly to moderately positively correlated with DSC for 7 raters (*ρ* = 0.15 to 0.50). All corresponding Adjacent:SC and DSC values, for each imaging slice of each dataset, are shown in **Supplementary Figure 2**.

**Table 2.** Spearman Correlations Between Rater agreement (DSC) with reference masks and Coefficient of Variation (CV) and Adjacent:SC Ratio across all participants and slices (n=467).

| Rater | DSC vs. CV |  | DSC vs. Adjacent:SC |  |
| --- | --- | --- | --- | --- |
| | Spearman's $\rho$ | $p$-value | Spearman's $\rho$ | $p$-value |
| A | -0.16 | 0.007 | 0.15 | 0.018 |
| B | -0.31 | < 0.001 | 0.45 | < 0.001 |
| C | -0.14 | 0.036 | 0.22 | < 0.001 |
| D | -0.26 | < 0.001 | 0.23 | < 0.001 |
| E | -0.30 | < 0.001 | 0.48 | < 0.001 |
| F | -0.39 | < 0.001 | 0.42 | < 0.001 |
| G | -0.50 | < 0.001 | 0.50 | < 0.001 |
| H | -0.30 | < 0.001 | 0.23 | < 0.001 |

Qualitatively, the registration of functional images to template space and the inverse registration of the template atlas to the functional image both showed visible differences in alignment (**Figure 3**). For example, in Dataset 2, shown with registrations informed by the reference and rater H, the PAM50 template masks of GM and WM are clearly not co-localized.

**Figure 3:**
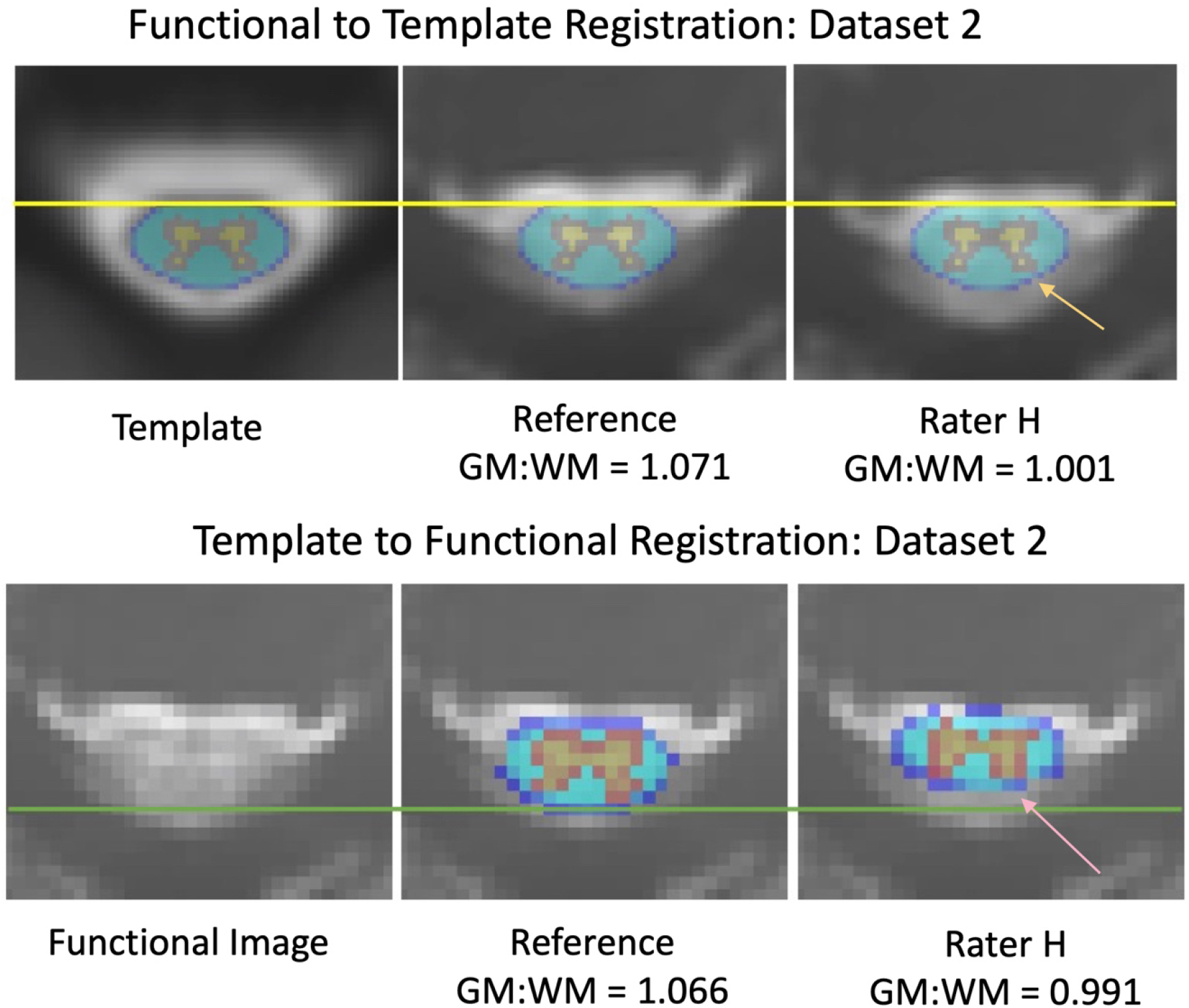
Differences in alignment between registration informed by reference and rater H masks, from dataset 2. The mean DSC was 0.736 across the axial slices for the input masks between the reference and rater for dataset 2. (Top) Registration of functional images to the PAM50 template. The yellow line represents the most anterior white matter voxel coordinate of the template mask. The orange arrow indicates an area of grey matter in the dorsal horn that is not aligned with the template mask in a registration informed by Rater H masking. (Bottom) Registration of the PAM50 template atlas to functional image space. The green line represents an estimate of the most dorsal coordinate of the functional image. The pink arrow indicates an area of grey matter in the dorsal horn that is not aligned with the atlas in Rater H registration. Accuracy of registration alignment is supported by the GM:WM ratio, as misalignment introduces a mixing of GM, WM, and potentially CSF voxels into the masked areas, reducing the observed contrast.

GM:WM contrast, calculated using the PAM50 template masks following spatial normalization of the functional data to the PAM50 template image (akin to the top panels in **Figure 3**), was greatest for the original study results (REF). **Figure 4** shows the GM:WM contrast results for all raters and datasets, maintaining the ordering of **Figure 2** with the REF results added on the top row. Across raters, this metric generally increased with higher agreement between RM and REF (with exceptions), as illustrated by a vertical gradient. As anticipated, there is also substantial variability of GM:WM contrast across the individual datasets, unrelated to rater masking and spatial normalization. ANOVA revealed that both rater and dataset had marked contributions to variance in GM:WM contrast, with dataset contributing relatively more variance, as anticipated (rater: *F*(8,160) = 23.57, *P* < 2×10^−16^, *η*^2^ = 0.24; dataset: *F*(20,160) = 22.00, *P* < 2×10^−16^, *η*^2^ = 0.56).

**Figure 4:**
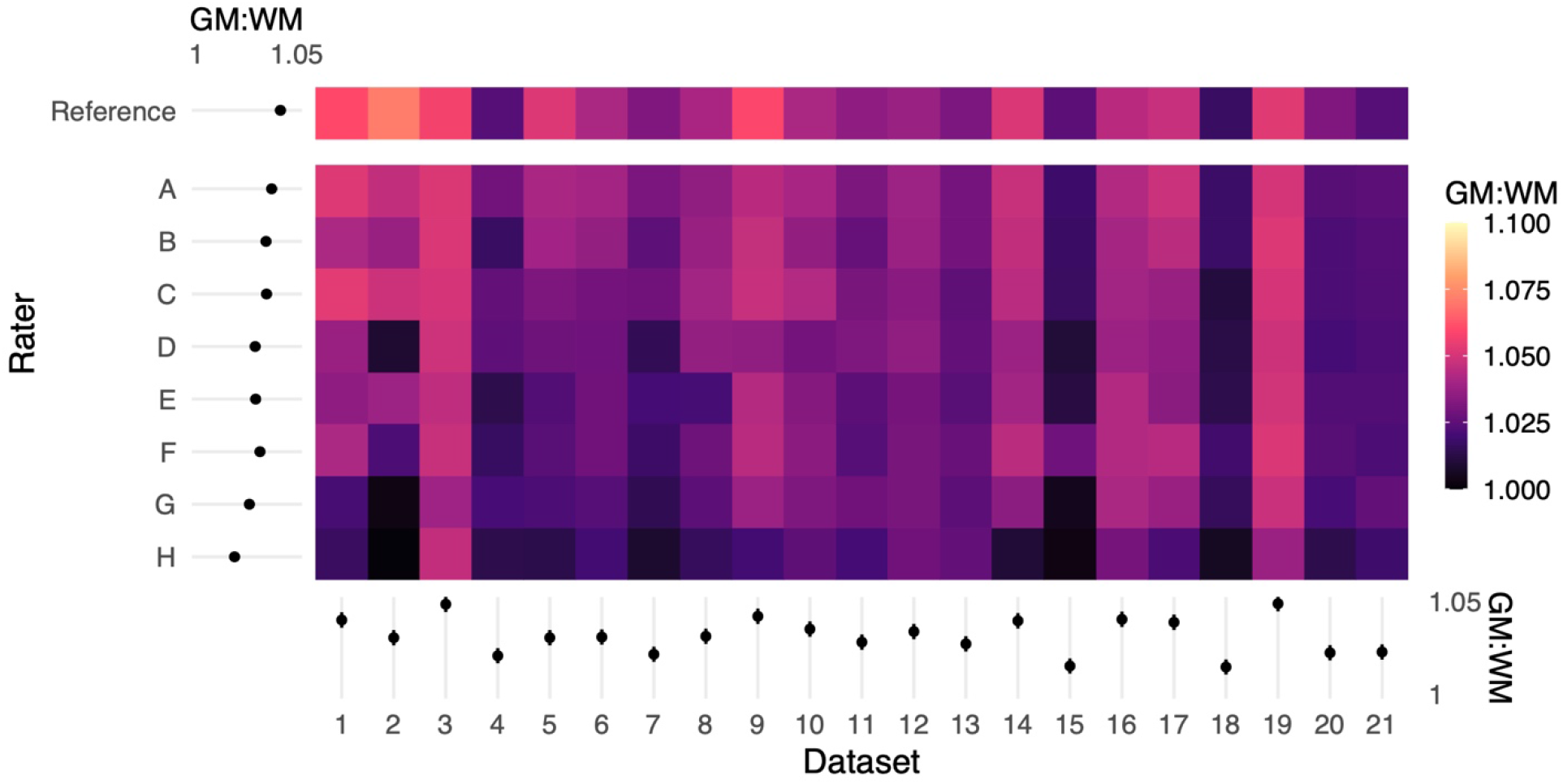
Color map of GM:WM contrast across all rater registrations. Higher GM:WM contrast for a given input dataset indicates relatively better registration alignment. The margins (left and bottom) contain estimated marginal means for each rater and dataset, and error bars indicate 95% CIs. The reference masks produced registrations with the highest GM:WM contrast. Rater mask agreement with the reference mask (logit-transformed DSC) is correlated with higher levels of GM:WM contrast following spatial registration. Moreover, there are marked dataset effects on GM:WM, which are reflected by the ANOVA results.

Fisher’s *z* spatial correlations between individual participant ipsilateral activation maps generated by each rater and the reference are shown in **Figure 5**. Horizontal and vertical trends are both present in the spatial correlations, indicating dataset and rater effects are both influencing the agreement of observed activation maps at the individual-level. Note, datasets were ordered by initial logit-transformed DSC agreement (as described for **Figure 2**) for this visualization.

**Figure 5:**
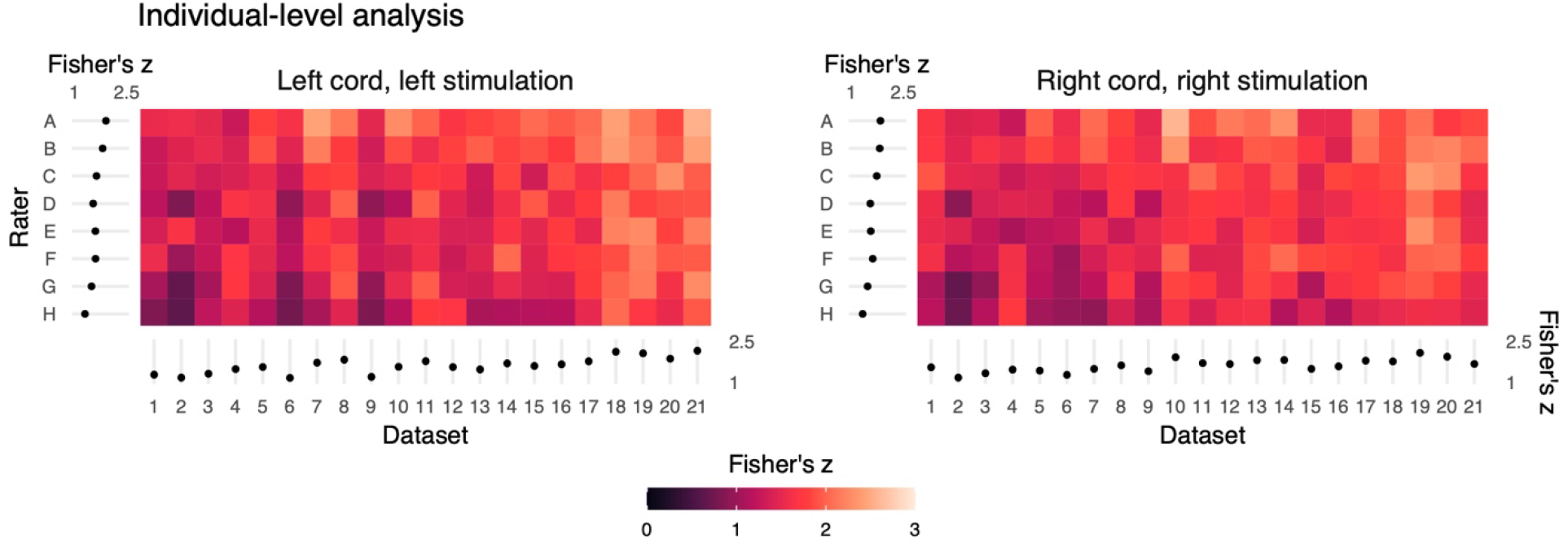
Summary of the individual-(or dataset-) level activation maps. Spatial correlation of statistical activation maps derived from the individual-level analysis results of each rater and the reference, for ipsilateral activation of left- and right-sided stimulation trials. Results shown as a color map of Fisher’s z correlations between rater and reference un-thresholded statistical activation maps, with the dataset order matching Figures 2 and 4. The horizontal and vertical gradient trends indicate that rater agreement with the reference mask and dataset factors influence agreement with the reference individual-level statistical activation maps. Marginal mean Fisher’s z scores by dataset and rater are shown at the axes. Error bars on the marginal means indicate 95% CIs.

Results from group-level analyses are also shown in **Figure 6** (left) and **Table 3**, where results from z-testing between the reference and raters for each stimulus and hemicord condition (*e.g*., Left hemicord, left stimulus) were distributed about 0 for all cases, illustrating no average Z-score differences.

**Figure 6:**
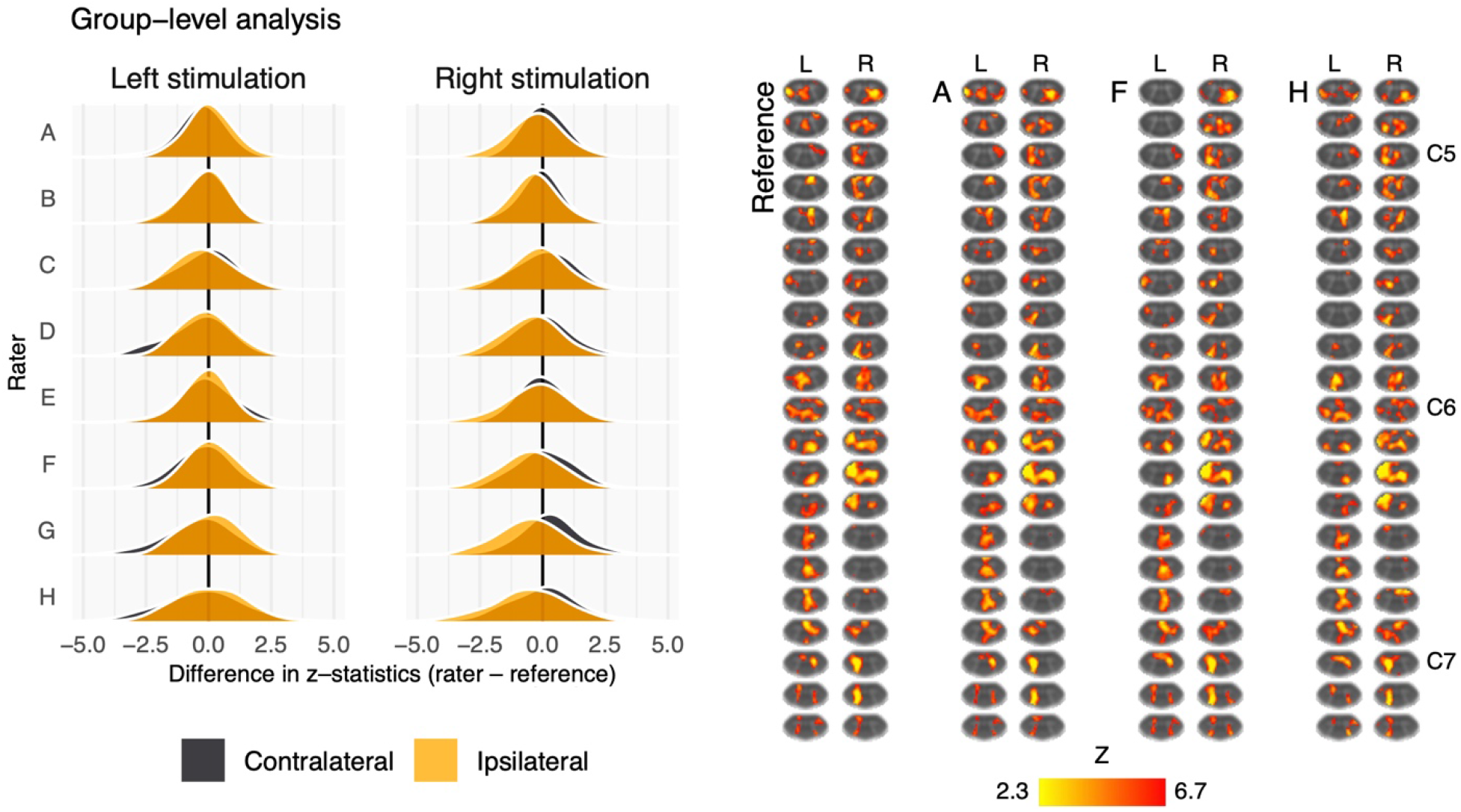
Summary of the group-level activation maps. Left: distribution of group-level activation z-statistics in the left and right hemicords for left- and right-sided stimulation trials. The shape and centering of the distributions are generally similar across all raters illustrating no systematic difference in average z-score. Right: group-level activation maps from the reference, rater A, rater F, and rater H. Activation is thresholded at Z-score>2.3 (cluster corrected p<0.05). Data have been transformed to the standard PAM50 template space, and the approximate level of C5-C7 spinal levels are indicated. (L=left, R=right sided stimulation).

**Table 3:** Magnitude of the reference’s group-level z-statistics relative to the raters’ distribution. The reference group-level z-statistic for each voxel was scaled by the distribution of z-statistics from the other raters. Presented here are the mean and standard deviations of the standardized reference z-statistics. A mean of 0 would indicate that the reference has the same z-statistic as the other raters (on average); values greater than zero indicate greater z-statistics than the raters’ average; and values lower than zero indicate lower z-statistics than the raters’ average.

| Side | Stimulation | Mean | ± SD |
| --- | --- | --- | --- |
| Left | Ipsilateral | 0.05 | $\pm 1.07$ |
| Right | Ipsilateral | 0.30 | $\pm 1.35$ |
| Left | Contralateral | -0.09 | $\pm 1.27$ |
| Right | Contralateral | 0.49 | $\pm 1.30$ |

Detailed in **Table 4**, spatial correlations between the raters’ and reference’s group-level results ranged from 0.954 to 0.875 (mean of 0.923) for the ipsilateral activations, and 0.952 to 0.781 (mean of 0.892) for the contralateral activations. Thus, while systematic trends were not observed across the results of different raters, there was observable disagreement in activation mapping at the group-level. An illustration of these differences is shown in **Figure 6** (right).

**Table 4:** Group-level activation map spatial correlations (Fisher’s z) between each rater and the reference by side (Left (L) or Right (R)) and stimuli condition (ipsilateral or contralateral activation).

| Rater | Side | Spatial Correlation<br>(Fisher's z) |  |
| --- | --- | --- | --- |
|  |  | Ipsilateral | Contralateral |
| **A** | *L* | 0.950 | 0.952 |
|  | *R* | 0.931 | 0.924 |
| **B** | *L* | 0.952 | 0.942 |
|  | *R* | 0.954 | 0.935 |
| **C** | *L* | 0.916 | 0.902 |
|  | *R* | 0.930 | 0.871 |
| **D** | *L* | 0.933 | 0.901 |
|  | *R* | 0.923 | 0.863 |
| **E** | *L* | 0.944 | 0.930 |
|  | *R* | 0.910 | 0.895 |
| **F** | *L* | 0.943 | 0.907 |
|  | *R* | 0.905 | 0.896 |
| **G** | *L* | 0.928 | 0.890 |
|  | *R* | 0.902 | 0.839 |
| **H** | *L* | 0.875 | 0.849 |
|  | *R* | 0.877 | 0.781 |

## Discussion

In this study, we assessed the potential impact of manual spinal cord contouring in native fMRI space, which is currently a recommended input to spatial normalization and group analyses in spinal cord fMRI. In a small cohort of 8 raters of varying experience, independently contouring the cord in 21 fMRI datasets acquired at 3T as part of a prior study, we observed mask differences attributable to both rater and dataset quality. Variability in masking was assessed by calculating DSC agreement with a reference rater (i.e., the rater from the original published work).

Regarding inter-rater variability, a priori expectation was that raters with more neuroimaging experience would achieve higher DSC agreement with the reference. While this was observed to be true at the extremes, there are notable deviations from this prediction: ordering raters by average logit-transformed DSC, one novice rater with no prior imaging experience (E) outperformed a researcher with 7 years of MRI experience (F), and a trainee with 1 year of spinal MRI experience (B) outperformed a researcher with 8 years of spinal MRI experience (C). Considering rater experience as categorized in Table 1, years of experience specifically in spinal cord fMRI (as opposed to neuroimaging or spinal cord MRI more generally) may be a more important factor. The obscured spinal cord boundary in fMRI data (due to partial-volume effects, low tissue contrast, and physiologic noise) may also lead to highly variable contouring performance among all non-experts. However, while rater F is surpassed by a novice rater (E) based on average logit-transformed DSC, they achieve a noticeably small spread of DSC values across all datasets. as visualized by **Supplementary Figure 1**. This suggests that more experienced raters may generate contours in a more consistent manner, therefore DSC with the reference may suffer as a result of systematic differences in interpreting the cord rather than inconsistent masking. Observations from downstream analysis suggest that this consistency in masking can lead to more robust downstream analysis: rater F overtakes raters D and E in GM:WM (**Figure 4**) and spatial correlation in individual-level analysis (**Figure 5**).

However, several limitations must also be acknowledged related to our choice of raters and their respective experience and training. First, while there is a wide range of MRI experience represented, all our raters have limited experience with spinal cord fMRI. We also acknowledge that there may be individuals more qualified than our reference rater to provide spinal cord masks for comparison (for example, a board certified neuroradiologist with equivalent years of spinal cord fMRI experience). We stress that the purpose of this study was to generate variability in masking by recruiting raters of varying levels of experience and describe how this variability affects the results of a previously published analysis pipeline. Our observations of downstream variability arising from this cohort of raters suggest that efforts should be made to standardize spinal cord masking to ensure robustness of results. One such way to increase robustness in masking may be to use the STAPLE method to combine segmentations from multiple experienced raters.[52] Another limitation is that the environment in which raters generated contours was not controlled. Therefore, although raters were instructed to take approximately 15 minutes per dataset, there may have been variability in the level of effort or focus put into the masking process, possibly confounding rater trends that were observed. Note that this effect is not expected to be systematic across datasets and should therefore not affect trends attributable to dataset.

Mask variability was attributable to dataset as well. Because these differences naturally occur along the edges of the cord, at the anatomical boundary of spinal cord white matter and surrounding CSF, we hypothesized that the signal contrast between these regions in each dataset would be associated with the agreement of rater masks with the reference. Indeed, better contrast at the edges of the cord (higher Adjacent:SC ratio) was significantly correlated with higher values of DSC, as shown in **Supplementary Figure 2**, which illustrates the slice-wise DSC and Adjacent:SC ratio correlations for every rater. Interestingly, we observe many datasets in this study presented with Adjacent:SC ratios less than 1, which is not anticipated given the T2* of tissue versus CSF. Furthermore, this ratio shows added variability across longitudinal image slices of each dataset. Signal dropout that could cause CSF voxels to appear darker than tissue voxels may be a result of susceptibility artifacts due to magnetic field inhomogeneity and intra-voxel dephasing through the use of thicker slices (3mm) in the functional acquisition. The breakdown of the expected positive contrast was often observed in the dorsal aspect of the cord, perhaps due to a posterior shift while a participant is being scanned in a supine position.[53] A posterior shift of the cord reduces the amount of CSF buffer between neural tissue and ligamentum flavum, disc, or bone from the spinal canal and may lead to increased partial volume averaging of these tissues. CSF flow may also impact the signal intensity of CSF voxels in this acquisition, potentially increasing or decreasing voxel brightness, thus influencing the Adjacent:SC ratio and mask fidelity. (Note, the effect of this contrast breakdown on rater masking was also captured by the negative association between CV of adjacent voxels and DSC: inconsistent brightness among voxels surrounding the cord led to decreased agreement of the rater mask with the reference mask.)

This study used the temporal mean functional image for contouring the spinal cord, which may merge these flow artifacts and reduce apparent image contrast. It may be more successful to contour the spinal cord on one fMRI volume, rather than the temporal mean image, if a volume with maximal Adjacent:SC contrast can be identified. However, it will be challenging to do this in a robust and systematic manner and appropriately integrate this step into volume realignment (i.e., motion correction) procedures, and this would not fully compensate for inherently poor Adjacent:SC contrast across the scan. These findings support the need for continued improvement in spinal cord fMRI acquisition techniques to improve and stabilize image contrast (and particularly tissue-CSF contrast) along the length of the cord while mitigating flow artifacts, such as improved receive coils[18,54-55], higher static magnetic field strengths[18,54], sequence and protocol optimization[18,41-42], and image processing techniques [14-19,54].

Following spatial normalization to a template image, the functional image GM:WM ratio (using GM and WM masks from the PAM50 template) was also reflective of both rater and image quality. In T2*-weighted images, GM is expected to be brighter than WM, and thus a well-registered functional dataset will yield a more positive GM:WM contrast compared to a less successfully registered version of the same input data. Indeed, registration informed by the reference masks achieved the highest GM:WM values in 15 of the 21 datasets. In our results (**Figure 4**), rater disagreement with the reference mask appeared associated with lower GM:WM ratios following image registration, suggesting that the underlying tissue projections onto the template contained a mix of tissue classes. These results demonstrate how a rater’s manual contouring of the spinal cord in native fMRI space influences the success of image registration to template space.

Reducing the dependency of the image registration algorithms on manual inputs could potentially mitigate many of these confounds: as has been done for the registration of high-resolution anatomical images of the spinal cord, convolutional neural networks could be trained for automated cord segmentation in fMRI datasets.[47] Such advances would have the added benefit of speeding up the image processing pipeline by removing the rate-limiting step of manual contouring, and would have the added benefit of improving analysis repeatability and facilitating robust sharing and combining of spinal cord fMRI data resources.

Differences in masking were also shown to impact the spatial distribution of activation at the individual-level, shown in **Figure 5**, however the impact on group-level results is less obvious (**Figure 6**). This may be due to averaging over a relatively large voxel size (1×1×3mm) when evaluating the cord with an average size of 7.4±0.9mm anterior-posterior and 11.4±1.2mm left-right at the C7 level.[56] The “straightening” of the cord inherent to registering individual spinal cord anatomy to the PAM50 template will produce subject-specific interpolation effects that could also influence the accuracy and sensitivity of group-level activation results. Additionally, the neural activation of interest, due to the tactile stimulus, is expected near the center of the cord, in GM, where rater masks are most likely to agree (**Supplementary Figure 3)** and the nonlinear registration algorithm may produce more consistent results. (Note that large BOLD signal changes may also occur in draining venous vessels, spanning both central and peripheral regions of the spinal cord.) **Figure 6** (right) illustrates the group-level activation maps achieved using reference and example rater masks. While some spatial differences are observable (for example, Rater F misses activation to the left stimulus in superior slices of the cord), differences in spatial correlations of unthresholded activation patterns (Table 3) do not result in large qualitative differences in thresholded activation maps.

To further investigate any systematic differences between the group-level activation maps for each rater and the reference rater, difference maps (**Supplemental Figure 4**, left) and a non-parametric one-sample t-test with threshold-free cluster enhancement were calculated. The Z-score difference map tends negative in some regions, such as C6, (i.e., raters’ activation is less than that of the reference). **Supplemental Figure 4** (right) shows voxels in which the difference between raters and the reference is significant (P<0.05, family-wise error rate controlled). Although this provides insight into the spatial distribution of activation differences between the raters and reference, it is a generous test and assumes the reference rater results to be a perfect ground-truth, as the variability of the reference rater masking and activation results is unknown.

Finally, the lack of robust, systematic differences in the group-level results (despite clear impact on individual-level results) may also simply reflect the inherent challenges of measuring BOLD responses in the spinal cord, where there are poorly resolved physiologic motion confounds and a small anatomical target relative to the image resolution. In the original work, the classical definitions of dermatomal sensory distributions were not clearly observed in the group-level activation to left and right sensory stimuli.[37] Although predominantly ipsilateral activations were observed, they were not localized to the dorsal aspects of the cord, and activation spread across vertebral levels rather than localizing to the expected C7 region. One interpretation is that there is a more complex anatomical network underlying spinal neurological response to stimuli, and we refer the reader to the original work for a more detailed neurophysiological interpretation. However, it could also suggest that there remains a fundamental obstacle of low signal-to-noise ratio in spinal cord fMRI that hinders robust mapping of true activations. The results of this study indicate that such limitations in spinal cord fMRI sensitivity and specificity may be more critical to activation mapping than the subtle variations in image co-registration that arise from manually contoured spinal cord masks. However, as image quality improves, through developments in hardware, acquisition strategies, and image processing techniques, it may become apparent that co-registration of functional data to a standard template space is increasingly important in achieving accurate and robust group-level results.

## Conclusion

We observed differences in individual rater masks of the spinal cord in fMRI data when compared to masks from a reference rater. These differences were driven by both rater and dataset effects, and led to variable co-registration with a standard spinal cord template image. This variability propagated into differences in individual-level fMRI activation results, as measured via spatial correlation between the reference and raters’ activation maps for left and right sensory stimuli. However, when performing group-level analyses, these masking and co-registration differences did not have a systematic effect on the average Z-score of resulting group-level activation. While increasing consistency in manual contouring of spinal cord fMRI data could improve data co-registration and ultimately the inter-rater agreement in activation mapping, our results suggest that other improvements in image acquisition and post-processing may be more critical to address. Automated approaches for segmenting the spinal cord in fMRI data, although potentially inferior to an expert manual segmentation, would speed processing times and potentially reduce rater bias in the analysis pipeline. Future work to ensure robust processing of functional imaging data is needed to improve the sensitivity and specificity to true neural activations in the human spinal cord.

## Acknowledgements

Research supported by a grant from the Craig H. Neilsen Foundation (595499), and by grants from the National Institute of Neurological Disorders and Stroke (K23NS091430, K23NS104211, and L30NS108301). K.J.H. was supported by an NIH-funded training program (T32EB025766). A.D.V. was supported by the National Science Foundation Graduate Research Fellowship (DGE-1324585) and by the National Institute of Neurological Disorders and Stroke of the National Institutes of Health (F31NS126012). The content is solely the responsibility of the authors and does not necessarily represent the official views of the National Institutes of Health.

## Appendix A

Spinal Cord Toolbox commands utilized to register functional images to the PAM50 template.

~~~
sct_deepseg_sc
  -i anatomical_image
  -c t2
  -centerline svm
  -kernel 2d
sct_register_to_template
  -i anatomical_image
  -s anatomical_image_segmented.nii.gz
  -l anatomical_image_vertebrae_labels
  -c t2
sct_register_multimodal
  -i PAM50_t2*_template_image
  -iseg PAM50_spinal_cord_mask
  -d functional_mean_image
  -dseg functional_mean_image_spinal_cord_mask
  -param
    step=1
       type=seg
       algo=centermass
    step=2
     type=seg
     algo=bsplinesyn
     slicewise=1
     iter=3
-initwarp template_to_anatomical_image
-initwarpinv anatomical_image_to_template
~~~

**Supplemental Figure 1:**
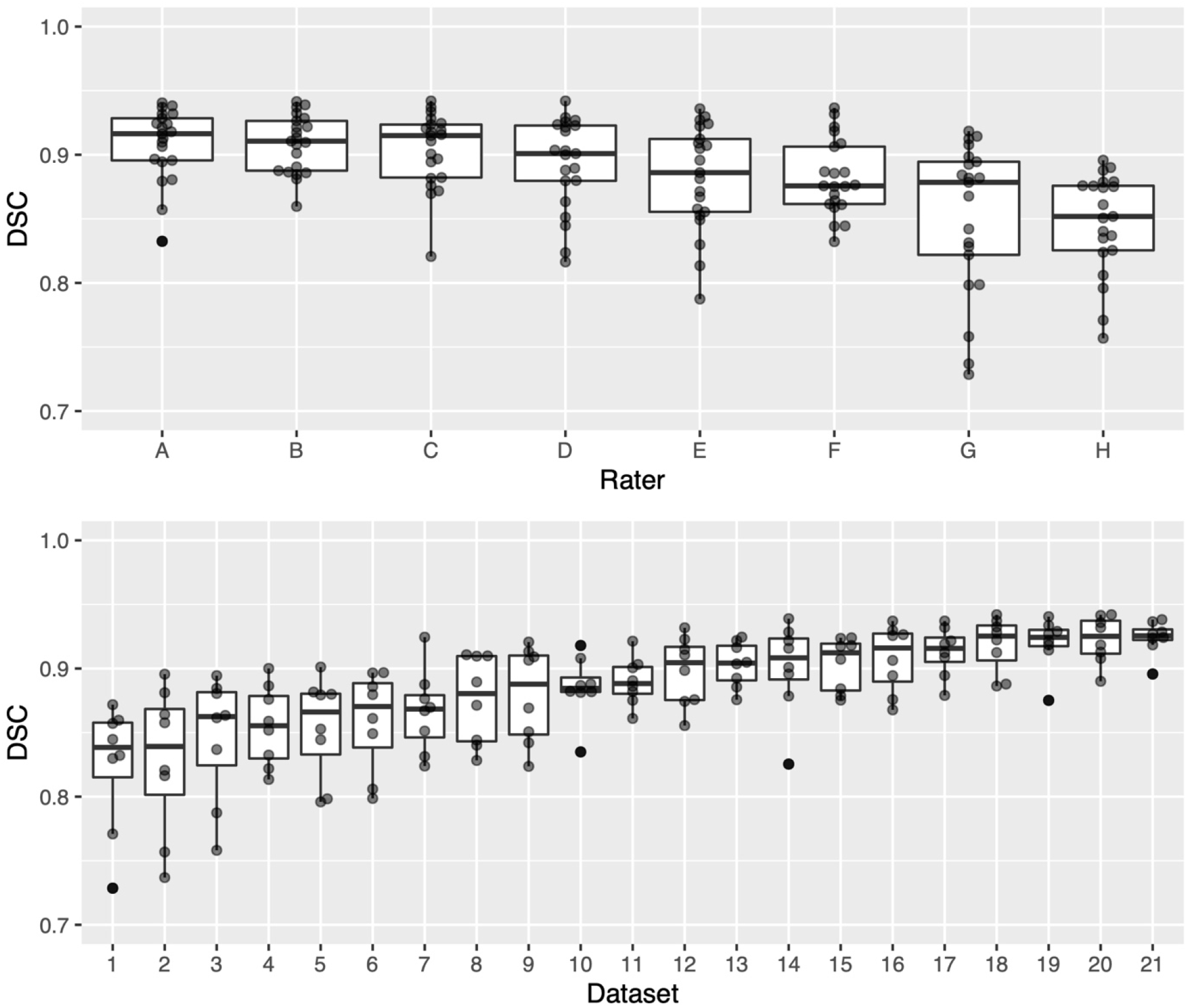
DSC of each rater mask compared to the reference mask visualized as box-and-whisker plots. Top: DSC plotted by rater. Bottom: DSC plotted by dataset.

**Supplemental Figure 2.**
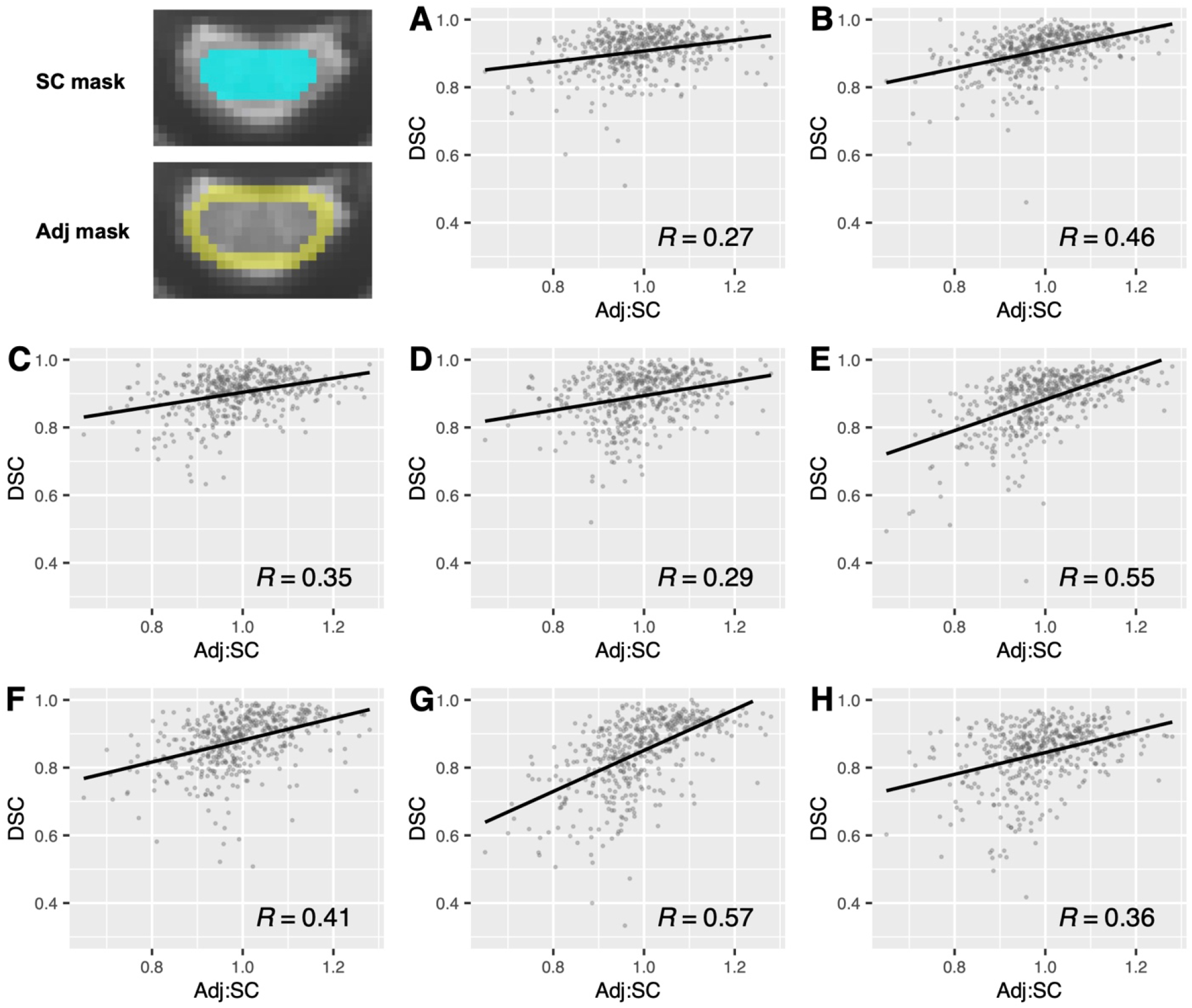
Slice-wise DSC and Adj:SC ratio correlations for every rater. Top left: a visualization of masks defining adjacent and SC voxel regions on the reference mask. Plots A-H: DSC and Adj:SC ratio are positively correlated for every rater. Greater contrast between spinal cord and adjacent voxels on a given transverse slice is associated with increased agreement with the reference (positive correlation).

**Supplemental Figure 3:**
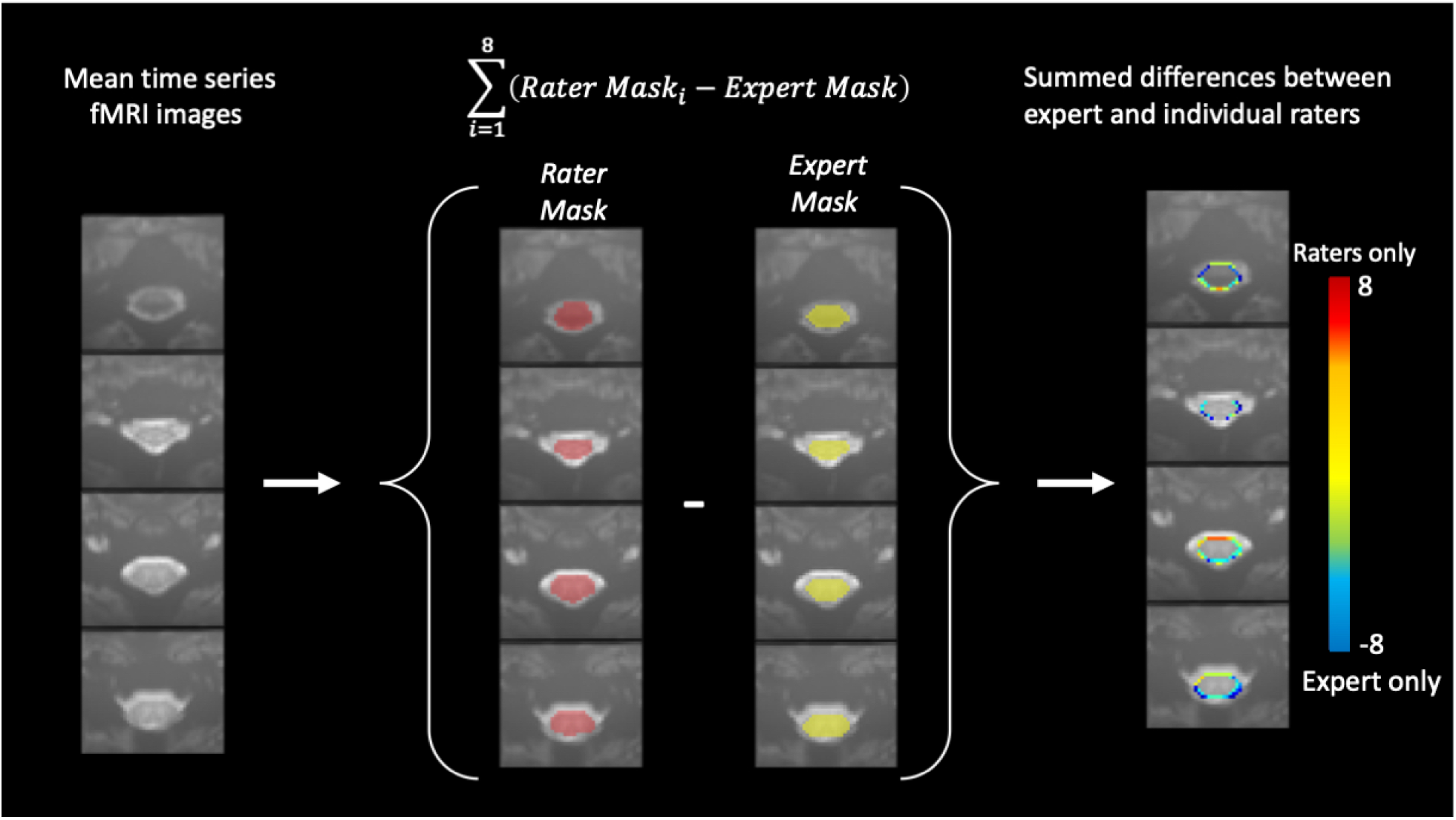
An example of the summed differences between the reference and the raters’ mask. Eight raters and reference rater contoured the spinal cord on temporal mean fMRI images. The reference mask was subtracted from each of the rater masks and the differences summed. The resultant summed difference maps are shown on the right. The difference map colorscale runs from -8 to +8, with negative values representing voxels included in the reference mask were not selected by the 8 raters, and positive values being the reverse, where the raters included voxels that were not selected by the reference. The difference maps illustrate the location of disagreement between the raters and reference is at the edges of the spinal cord.

**Supplemental Figure 4.**
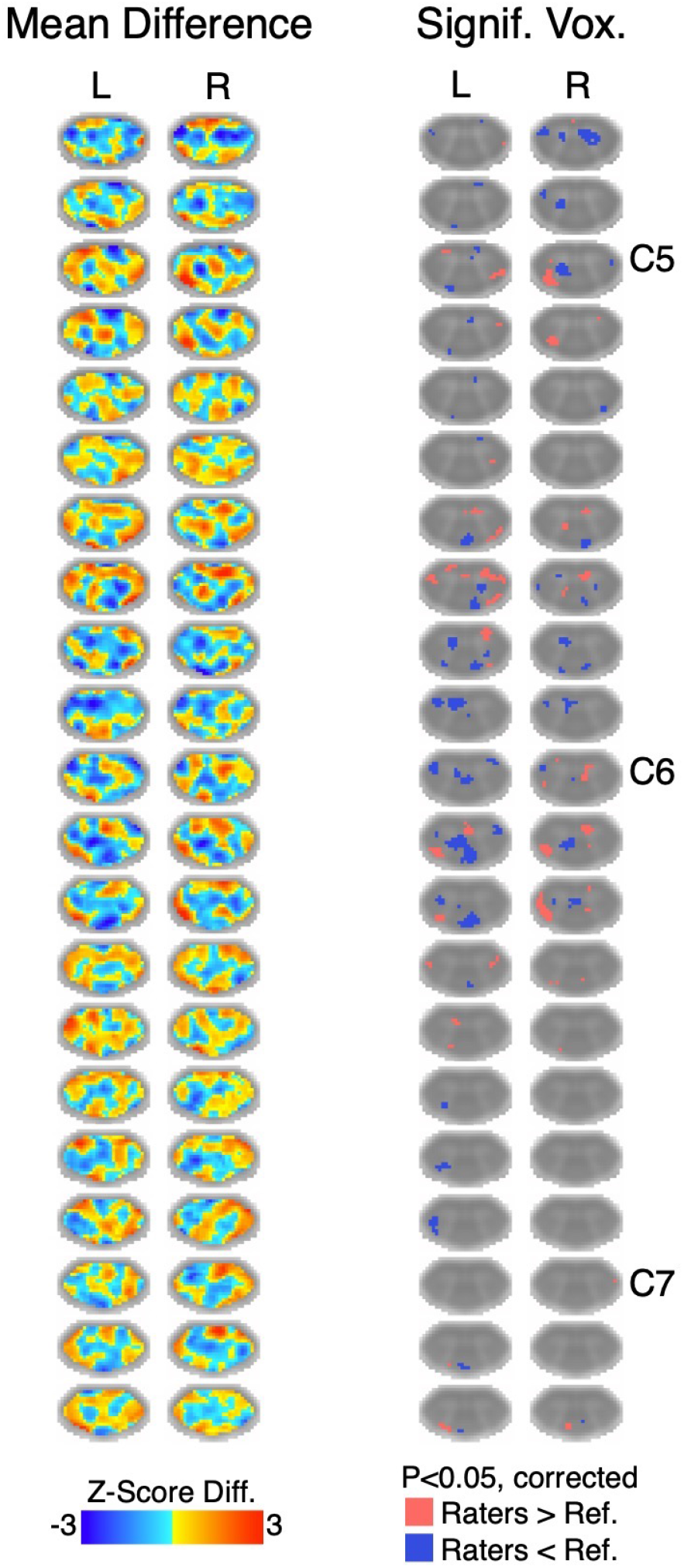
Left: mean difference of group-level activation Z-score maps (shown in Figure 6) between the reference and each of the 8 raters (e.g., rater A - Reference) for the left and right contrasts. Right: map of significant voxels from non-parametric one-sample t-test with threshold-free cluster enhancement (P<0.05, family-wise error rate controlled). For the left contrast, 6.20% (2.06% Raters > Ref., 4.14% Raters < Ref.) of voxels in the spinal cord were significantly different between the 8 raters and the reference rater. For the right contrast, 4.25% (1.90% Raters > Ref., 2.35% Raters < Ref.) of voxels in the spinal cord were significantly different between the 8 raters and the reference rater. Slices shown are the same as in Figure 6.

